# Chilean blind spots in soil biodiversity and ecosystem function research

**DOI:** 10.1101/2021.06.24.449754

**Authors:** César Marín, Javiera Rubio, Roberto Godoy

## Abstract

Soil harbor up to a quarter of the world’s biodiversity, contributing to many ecosystem functions. It is of great importance to identify distribution patterns of soil organisms and their ecosystem functions to support their conservation and policy building. This has been recently analyzed at macroecological scales, but analyses at national/local scales are scarce. Here we identify and analyze the blind spots in soil taxa and ecosystem functioning data in continental Chile, through a Web of Science articles (1945-2020) search, and focusing on ten soil taxonomic groups and four ecosystem functions (nutrient cycling, decomposition, water infiltration, soil respiration). A total of 741 sampling sites were obtained from 239 articles. In 49.25% of the sites soil biodiversity was studied, while this percentage was 32.65% for ecosystem functions; in 18.10% of the sites both soil biodiversity and ecosystem functions were investigated at the same time, a surprisingly high percentage compared to global studies. By far, Bacteria/Fungi and nutrient cycling were the most investigated taxa and function, respectively. There is a significant number of soil taxa (Acari, Collembola, Nematoda, Formicoidea, Protista, Rotifera) represented by just a few sites concentrated in specific Chilean regions. Places like the central regions, the Atacama desert, and the Valdivian temperate forests present a proliferation of studies on soil Fungi, Bacteria, and nutrient cycling, reflecting historical interests of established research groups. Based on this research, we are identifying the causes of the data blind spots and invite the Chilean soil ecology community to propose ideas on how to fill them.

## Introduction

Soil is a highly diverse habitat which contains a plethora of very different organisms, ranging from bacteria and fungi, to nematodes, earthworms, and moles, among others. It is estimated that soil harbor up to a quarter of all living species on Earth, and that one gram of healthy garden soil may contain one billion of bacterial cells, up to one million individual fungi, about one million cells of protists, and several hundreds nematodes (European Commission 2010). Soil biodiversity is a main actor driving several ecosystem functions and services, including nutrient cycling, regulation, and plant acquisition, plant productivity, reduction of plant pathogens, control of antibiotic resistance genes, climate regulation, food production (Bardgett and van der Putten 2014; Delgado-Baquerizo et al. 2020), etc.

Despite its importance, soil biodiversity has historically and largely been neglected. But progress has been made in recent years, for example with the launching of the first global report on the *State of knowledge of soil biodiversity* (FAO et al. 2020), which included contributions of more than 300 scientists worldwide. Nevertheless, one of the main questions in soil ecology still is how to causally relate soil biodiversity and ecosystem functioning at different spatiotemporal scales (Eisenhauer et al. 2017). Although over the last decade, global-scale studies have started to disentangle this causal relationship (Maestre et al. 2012, 2015; Pärtel et al. 2016; Delgado-Baquerizo et al. 2016, 2017, 2020; Soliveres et al. 2016; Song et al. 2017; Crowther et al. 2019), there is much work to be done compared to aboveground ecosystems, where for example the relationships between plant community attributes and productivity are well established (Flynn et al. 2011; Grace et al. 2016; Liang et al. 2016; Duffy et al. 2017). Still, it is important to establish causal paths between soil microbial communities attributes and ecosystem functions/services (Xu et al. 2020). Hall et al. (2018) defines three categories of microbial communities attributes: microbial processes (ie. nitrogen fixation), microbial community properties (ie. biomass C:N ratio, functional gene abundance), and microbial membership (ie. taxonomic and phylogenetic diversity, community structure, co-occurrence networks). In this conceptualization, microbial processes more directly affect a nutrient pool or flux, while the effects of community properties and microbial membership are more indirect, mediated by their concatenate effect on microbial processes.

Over the last decade there is a paramount of global soil ecology studies focusing on bacteria (Delgado-Baquerizo et al. 2018; Cano-Díaz et al. 2020), protists (Singer et al. 2019; Oliverio et al. 2020), fungi (Tedersoo et al. 2014; Egidi et al. 2019; Větrovský et al. 2020) including mycorrhizal fungi (Davison et al. 2015; Soudzilovskaia et al. 2015), invertebrates in general (Bastida et al. 2019), nematodes (van den Hoogen et al. 2019), earthworms (Briones and Schmidt 2017; Phillips et al. 2019), isopods (Sfenthourakis and Hornung 2018), ants (Gibb et al. 2017; Bertelsmeier et al. 2017), termites (Buczkowski and Bertelsmeier 2016), roots (Iversen et al. 2017), and the overall soil community (Fierer et al. 2009; Bahram et al. 2018; Cameron et al. 2019; Crowther et al. 2019; Delgado-Baquerizo et al. 2020; Guerra et al. 2020; Johnston and Sibly 2020; Luan et al. 2020). Despite these great advances in global soil ecology, major taxonomic, functional, geographic, and temporal gaps still exist (Bueno et al. 2017; Cameron et al. 2019; Guerra et al. 2020). Filling these gaps is crucial for soil biodiversity conservation and governance. Furthermore, and unlike aboveground biodiversity, there is no monitoring of soil biodiversity; thus, as much of global soil biodiversity is yet to be described, we do not even know at what pace are these unknown species being lost. There is urgent need for action.

Recently, when analyzing 17,186 sampling sites at a global scale (from macro-ecological scale studies) Guerra et al. (2020) found that just in 0.3% of those sites, both soil biodiversity and ecosystem functions where investigated at the same time. As both are so interdependent, much work needs to be done in order to integrate conceptually, causally, and disciplinary (regarding different knowledge areas) soil biodiversity and its associated ecosystem functions and services. Guerra et al. (2020) found a lack of conjoint studies of soil biodiversity and ecosystem functions at continental and global-scale studies, but, would the same happen at national, regional, or local scales? To find out, we conducted a similar analysis (searching Web of Science articles, extracting their coordinates, and assigning each site to the soil taxa and/or function investigated) restricted to the continental Chilean territory. Chile is the longest country in the World, and as such, it contains varied ecosystems: the driest desert (Atacama), high Andean ecosystems, Mediterranean climate areas, extremely rainy temperate forests, and Patagonian forests and steppe. There is plenty of interest in the soil microbial biodiversity of the country, reflected for example in the creation of The Chilean Network of Microbial Culture Collections (Santos et al. 2016), or more specifically with the Atacama (microbiome) Database (Contador et al. 2020).

This study aimed to identify the information blind spots in soil taxa and ecosystem functions in the continental Chilean territory, analyzing their distribution patterns, as a base line for future monitoring and conservation initiatives.

## Materials and Methods

### Literature search and coordinates extraction

In January 2021, a Web of Science search of articles published between 1945 to 2020 was conducted focusing on 10 soil taxa (Bacteria, Fungi, Archaea, Oligochaeta, Acari, Collembola, Nematoda, Formicoidea, Protista, Rotifera) and four ecosystem functions (nutrient cycling, decomposition, water infiltration, soil respiration) according to Guerra et al. (2020). The following keywords were used: (Chile* OR Arica OR Parinacota OR Tarapacá OR Valparaíso OR O’Higgins OR Maule OR Ñuble OR Biobío OR Araucanía OR Aysén OR Magallanes OR Metropolitan OR Antofagasta OR (Northern AND Chile) OR (Central AND Chile) OR (Southern AND Chile) AND (soil* OR belowground) AND (*function* OR *diversity OR organism* OR biota OR animal* OR invert* OR fauna*) AND distribution AND (*mycorrhizal* OR microb* OR nematodos* OR bacteria* OR ant* OR fung* OR invertebrate* OR earthworm* OR protist* OR eukaryot* OR collembola* OR rotifer* OR Archaea OR formic* OR mite* OR termite* OR arthropod* OR respiration OR decomposition OR nitrogen-cycling OR nutrient cycling OR water infiltration OR aggregate* OR bioturbation).

This variety of keywords were used in order to capture the maximum number of published articles, which often used very different expressions when referring to the Chilean administrative regionse, soil taxa, and ecosystem functions. Words like “Los Ríos” and “Los Lagos” referring to the administrative regions of Chile named that way, were excluded, as searching them leads to studies conducted in rivers and lakes. A second, more detailed search was necessary for focusing on specific geographic regions, soil, and the taxa or function of interest, using for example the following keywords: (Chile* OR Arica OR Parinacota OR Tarapacá OR Valparaíso OR O’Higgins OR Maule OR Ñuble OR Biobío OR Araucanía OR Aysén OR Magallanes OR Metropolitan OR Antofagasta OR (Northern AND Chile) OR (Southern AND Chile) OR atacama) AND soil* AND mycorrhizal*).

Each article was checked individually, discarding those that did not imply soil extraction from continental Chile and that did not analyze at least one of the soil taxonomic groups or ecosystem functions defined. After compiling the articles, a database including coordinates (UTM system), citation, DOI identifier, and soil taxa and ecosystem function investigated was constructed in an Excel file, available at: https://figshare.com/s/c7b6dce6b12edfbc5e7d (DOI: 10.6084/m9.figshare.14838804).

### Spatial data processing and analyses

Data was georeferenced using Qgis 3.6 (QGIS.org. 2021) to create three point layers projected in WGS84. These were used to elaborate 4 spatial distribution representation cartographies, also using a shape layer of regional administrative boundaries, extracted from the “Infraestructura de datos Geoespaciales de Chile (IDE)” database (http://www.geoportal.cl/visorgeoportal/) and a shape layer of ecoregions extracted from the RESOLVE Ecoregions dataset (https://ecoregions.appspot.com/).

The first cartography used three shape layers: the first one related to sampling sites dealing with soil biodiversity, the second one for sites dealing with soil ecosystem functions, and the third one for sites dealing with both. For each layer, the parameters of points grouping (or cluster) were applied in Qgis 3.6 properties, assigning a tolerance distance of 50 km. For the second and third cartographies, the same tools and parameters were used, but applying 10 soil taxa shape layers (second cartography) and 4 soil ecosystem function shape layers (third cartography). For the fourth cartography the same three shape layers from the first cartography were used. For each one, the Qgis 3.6 tool Heatmap (Core Density Estimation) was used, applying a 2 km radius to cover concentrations within that range. The color gradient was adapted to the design of previous cartographies, categorizing from a low density (only 1 point), to a high density meaning the existence of over 10 sampling points. The RESOLVE Ecoregions shape layer (for continental Chile) was superimposed in this fourth cartography. All cartographies were projected in WGS84 / EPSG: 4326.

To analyze the representativeness (percentage and no. of samples) of the 10 soil taxonomic groups and of the 4 ecosystem functions in the ecoregions, the ecoregions layer, the 10 taxonomic groups layers, and the 4 ecosystem function layers were transformed to raster files with a 2 km resolution, assuming each sample point equals one pixel. For the ecoregions layer, a value of 1 to 7 was assigned to each pixel depending on which ecoregion it corresponds to (ie. pixels with a value of 1 correspond to the Atacama desert; pixels with a value of 2 correspond to the Central Andean dry puna). For point layers, a value of 10 was given to each point. Using a raster calculator, the values were processed by multiplying: Ecoregions raster * Point layer raster for each taxon and function. The result was 10 raster layers for taxa and 4 raster layers for functions with values from 1 to 7 and 10 to 70. Following the same example: all values of 10 correspond to the sampling points located in the Atacama Desert, and all values of 20 correspond to the sampling points located in the Central Andean dry puna. Using the Unique values report raster tool, a report was obtained for each layer showing the number of pixels for each value. These data were extracted and arranged in four Excel tables: two referring to the number of pixels for taxa and functions, and the remaining referred to the percentage of representativeness of each taxa and function, according to each ecoregion.

## Results

A total of 239 Web of Science articles were obtained for continental Chile, from which 111 deal with soil biodiversity, 89 deal with soil ecosystem functions, and 39 investigated both. From these articles, 741 sampling points were obtained (Fig. 1), showing a greater number of soil biodiversity sites in the administrative regions of Antofagasta (north) and Los Ríos (south) (Fig. 1a) and centered on Bacteria and Fungi (Fig. S1). The Andean part of the Coquimbo region, and the regions of Aysén and Magallanes showed the least soil biodiversity sampling points (Fig. 1a), while taxa like Formicoidea, Protista, and Rotifera did not surpass five studies. The central zone of Chile, and the regions of Antofagasta and Los Lagos showed a major number of soil ecosystem functions sites (Fig. 1b), with nutrient cycling being the most studied function with 300 sampling sites, while the remaining functions did not surpass 50 sampling sites (Fig. 1e). The La Araucanía region was the one where soil biodiversity and ecosystem functions were most conjointly studied (Fig. 1c).

**Figure 1.**
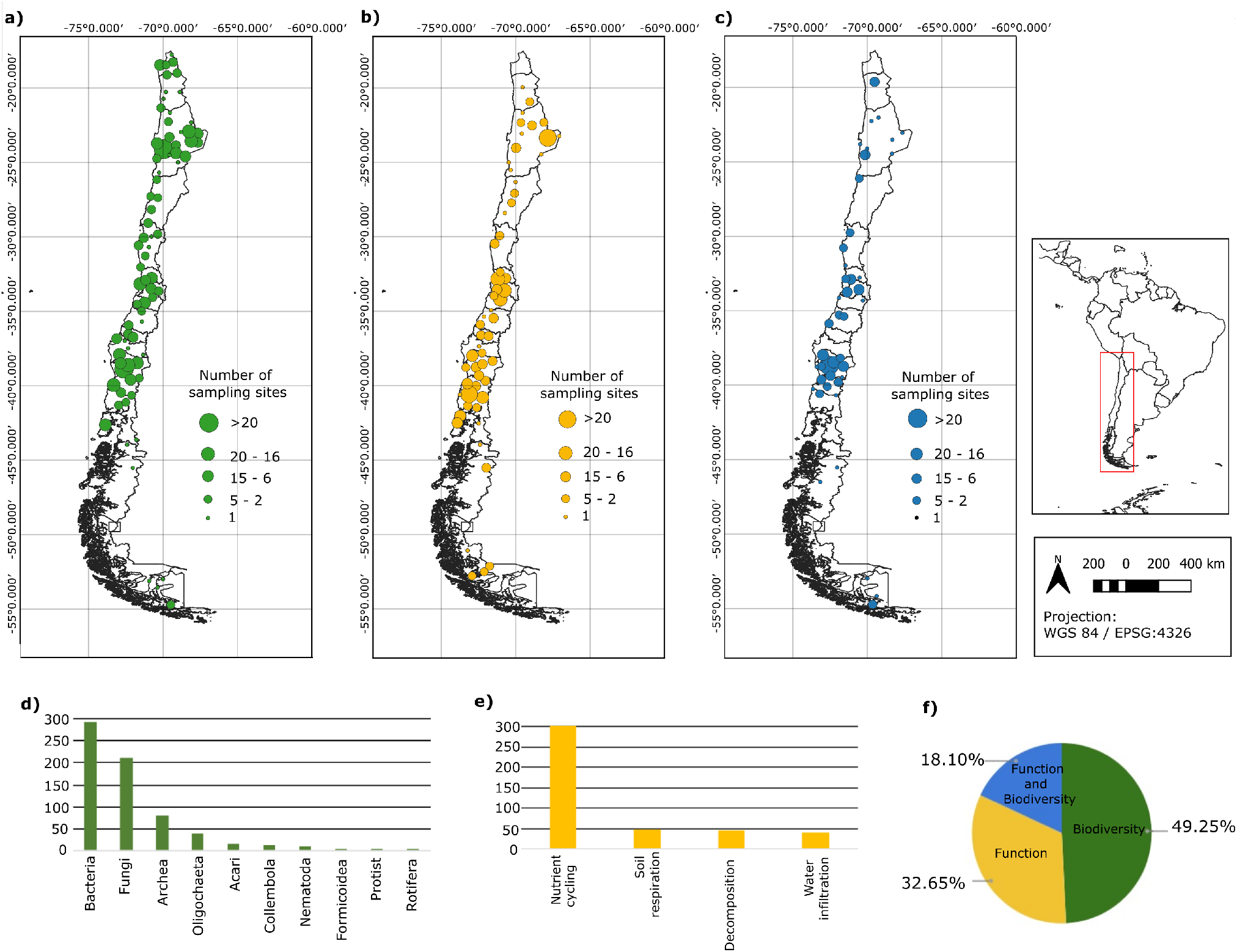
Distribution of sampling sites for soil taxa and ecosystem functions in continental Chile. **a.** Soil biodiversity sampling sites. **b.** Soil ecosystem functions sampling sites. **c.** Sampling sites where both soil biodiversity and ecosystem functions were conjointly studied. **d.** Number of sampling sites per soil taxa. **e.** Number of sampling sites per soil ecosystem function. **f.** Percentages of sampling sites investigating soil biodiversity, ecosystem function, and both. The size of the circles is based on a 50 km grid.

When doing a 2 km radius heatmap analysis, two hot spots in soil biodiversity sites were found: one at the south of the Atacama desert and one in the Chilean Matorral (which is also a hot spot for soil biodiversity and ecosystem functions when studied together; Fig. 2c), while some parts of the Central Andean dry puna presented medium density (Fig. 2a); this ecoregion also showed a hot spot for soil ecosystem functions sites (Fig. 2b). The Valdivian temperate forests had medium density regarding soil ecosystem functions sites (Fig. 2b). Ecoregions like the Magallanic subpolar forests and the Patagonian steppe had the highest sampling gaps (Fig. 2).

**Figure 2.**
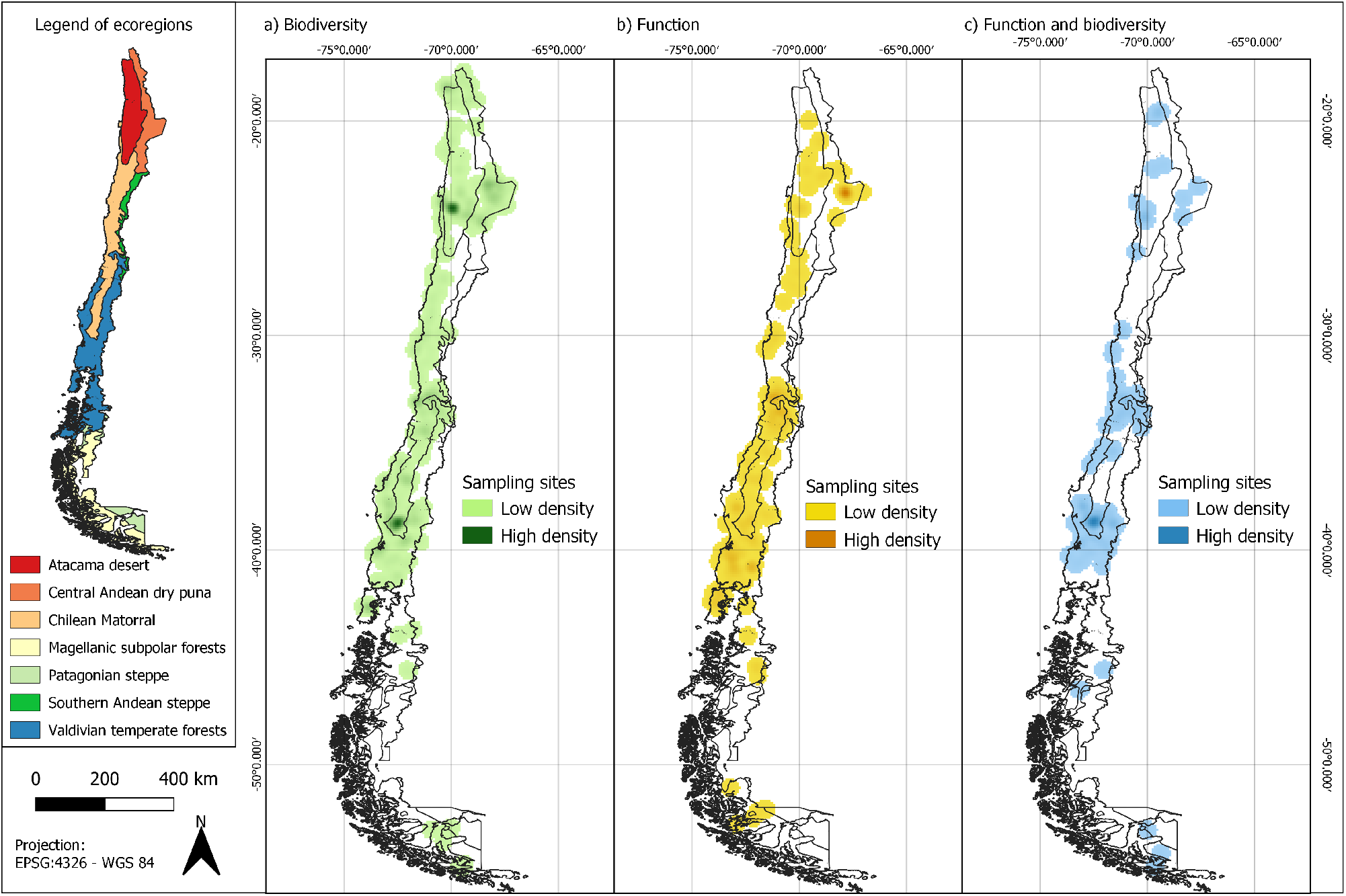
Heatmap of sampling distribution across continental Chile ecoregions (2 km grid). **a.** Soil biodiversity sampling sites. **b.** Soil ecosystem functions sampling sites. **c.** Soil biodiversity and ecosystem function sampling sites.

Regarding the representativeness of the 10 soil taxa and of the 4 ecosystem functions in the ecoregions, it was found that in all ecoregions, at least 5 soil taxonomic groups had a percentage coverage of less than 5% (Fig. 3a), with less than 10 sampling sites (Fig. 3b). For soil ecosystem functions, overall, a greater variability in percentage coverage (Fig. 3a) and number of sampling sites (Fig. 3b) was found. The number of sampling sites for soil ecosystem functions was generally low in at least five of the seven ecoregions, and did not surpass five sites for two ecosystem functions (Fig. 3b). Ecoregions like the Chilean Matorral and the Valdivian temperate forests had the highest soil ecosystem functions representativeness of number of sampling sites and coverage percentage (Fig. 3). The Magellanic subpolar forests, the Patagonian steppe, and the Southern Andean steppe presented extremely low numbers of sampling sites (Fig. 3b).

**Figure 3.**
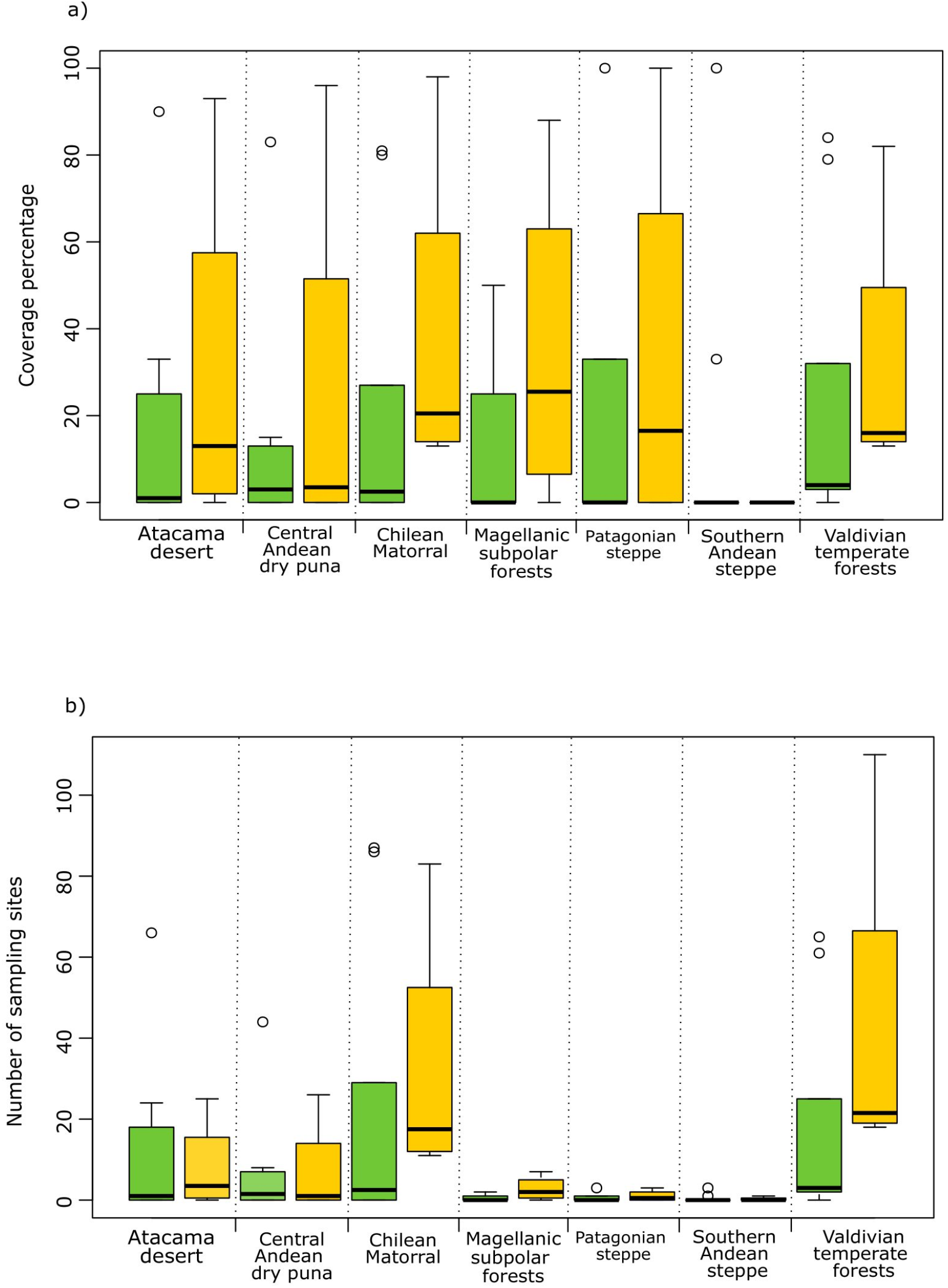
Representativeness of percentage coverage (**a**) and number of sampling sites (**b**) for soil biodiversity (green bars) and ecosystem functions (yellow) in the seven continental Chilean ecoregions.

Finally, historically there is an steady increase in Chilean studies dealing with soil biodiversity, ecosystem functions, and both (Fig. 4), especially during the last decade.

**Figure 4.**
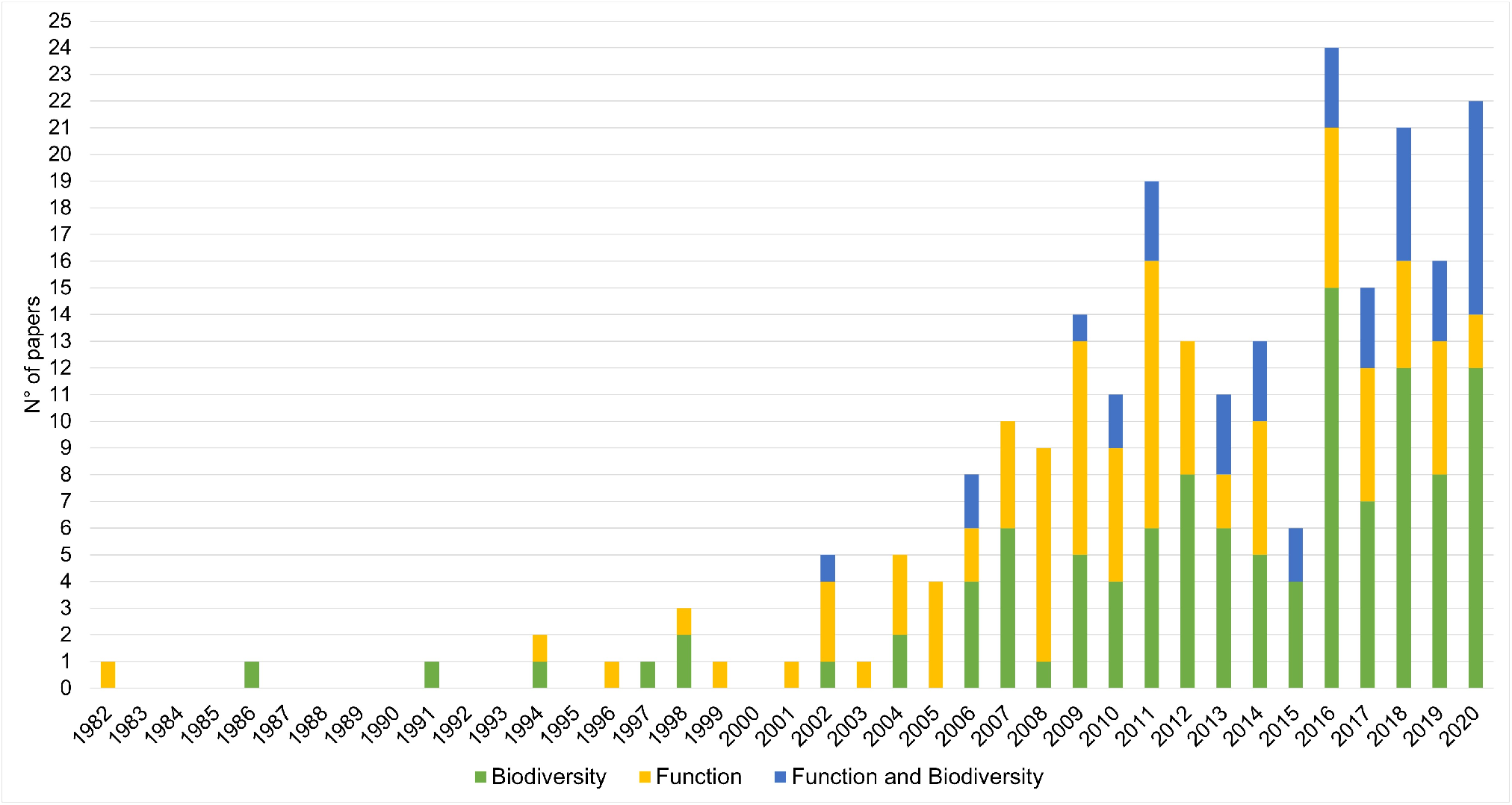
Number of Web of Science articles (N° of papers) published for continental Chile and dealing with soil biodiversity (green), ecosystem functions (yellow), and both investigated together (blue). The Web of Science search was for the period 1945-2020, but the first study appears in 1982.

## Discussion

In our analyses of soil biodiversity and ecosystem function research in Chile we overall found several types of biases: geographic, towards the Atacama desert, the central zone of Chile, and the Valdivian temperate forests; taxonomic, towards Bacteria and Fungi; and functional, towards nutrient cycling. Over the last decades, the Atacama desert, given its extreme conditions, has attracted plenty of national and international researchers interested in studying the microbial life under such conditions. So much so, that an special issue on the microbiology of the Atacama desert was launched by the journal Antonie van Leeuwenhoek in 2018 (Bull et al. 2018). Dry tephra of Atacama volcanoes (above 6000 m.a.s.l.) is the closest thing to the surface of Mars, as these “soils” are extremely acidic, oligotrhophic, and exposed to a thin atmosphere, high UV fluxes, and high temperate fluctuations (Schmidt et al. 2018). These conditions are perfect for the field of astrobiology, which also has proliferated in the Atacama desert. There are now important established Chilean research groups studying the Atacama soils microbial life. The central zone of Chile, around the Metropolitan and the Valparaíso regions, concentrate most of the population (52%), the most traditional and prominent universities, and the most crop productive area. This partially could explain the concentration of sampling sites around that zone. Finally, there was also a significant number of sampling sites (for soil biodiversity, ecosystem functions, and both) around the regions of La Araucanía, Los Ríos, and Los Lagos, in the Valdivian temperate forests. This reflects an historical interest of established research groups over the last four decades, as well as international collaborations, mainly originating from the Austral University of Chile and from La Frontera University (Godoy and Mayr 1989; Rubio et al. 1990; Godoy and Marín 2019).

Some soil taxa like Acari, Collembola, Nematoda, Formicoidea, Protista, Rotifera where barely studied, represented by just a few sites concentrated in some administrative regions. We think several reasons could explain this: i) An historical lack of interest in such groups, as for example the established research groups mentioned above have focused in Bacteria (for the Atacama desert) and Fungi (for Valdivian temperate forests); ii) The difficulties that the sampling for some of those groups carry out; iii) The relatively simple methods for Bacteria, Fungi, and Archaea sampling from soil, especially over the last decade with next generation sequencing techniques; and iv) All of the above combined. This trend where soil Bacteria and Fungi are the most studied taxa is not unique to this study (Guerra et al. 2020), and besides reflecting the ubiquity of such taxa (Tedersoo et al. 2014; Delgado-Baquerizo et al. 2018; Egidi et al. 2019; Cano-Díaz et al. 2020; Větrovský et al. 2020), it also shows their central role in ecosystem functioning, as usually taught in soil ecology. Perhaps the other soil taxa should be more investigated to disentangle unknown relationships with ecosystem functions. Also, as “nutrient cycling” encompass a great number of processes, it is understandable that this was the soil ecosystem function with most sampling sites.

Ecoregions like the Magellanic subpolar forests, the Patagonian steppe, and the Southern Andean steppe are of extremely hard access, with few to any populated center and University or research center nearby. These regions presented very few sampling points for soil biodiversity and ecosystem functions in our study. Despite this, they are very interesting from an aboveground perspective, showing high plant richness (for the Magellanic subpolar forests; Rozzi et al. 2008) and complex biodiversity patterns across geographic zones and vegetation types (for the Patagonian steppe and the Southern Andean steppe; Peri et al. 2016). Ideally, all ecoregions of Chile in all their extension should have at least medium density of sampling sites dealing with soil biodiversity and ecosystem functions at the same time, which is far from being the case; thus, there is plenty of work to be done.

In the 2019 United Nations Climate Change Conference (COP 25), Chile (co-organizer) presented an unprecedented upgrade on the state of its biodiversity, particularly recommending improving, strengthening, and implementing soil biodiversity monitoring programs, being soil one of the most vulnerable ecosystem components (Rojas et al. 2019). It was emphasized that an intensified soil use, inadequate agricultural practices, grazing, agro-forestry, and urbanization lead to well-known soil threats like erosion, pollution, acidification, nutrient leaching, salinization, loss of biodiversity and organic matter, among many others (Rojas et al. 2019). Also during COP 25, the main causes of native biodiversity loss where identified (for the period 1995-2016; Marquet et al. 2019): for the regions of Valparaíso, Metropolitan, O’Higgins, Los Lagos, and Magallanes it was the replacement of natural ecosystems for meadows and shrubs, previous to livestock and urbanization processes. For regions like Maule, Biobío, La Araucanía, and Los Ríos, the main cause of biodiversity loss was the replacement of native forests for commercial, fast-growing plantations (Marquet et al. 2019), like *Pinus radiata*, which retains high amounts of nitrogen, negatively affecting soil biodiversity (Oyarzún et al. 2007).

Now with a clearer picture of which are the geographic (all regions of Chile except for the central zone, and parts of the Atacama desert and the Valdivian temperate forests), taxonomic (all soil taxa except for Bacteria and Fungi, on the above-mentioned regions), and functional (all soil ecosystem functions except for nutrient cycling, on the above-mentioned regions) gaps of soil ecology in Chile, we can call for action. First, we need to feed the database constructed on this study, so if maybe some studies dealing with soil biodiversity and/or ecosystem functions in continental Chile were not included, please contact us. Second, as a Chilean soil ecology community there is a need for open data and open collaborations. Third, we need even more integration between the researchers investing soil biodiversity and those dealing with soil ecosystem functions. And fourth, we need specific legislation for soil biodiversity *per sé* (besides its importance for ecosystem functioning and food production), for conserving such biodiversity (Guerra et al. 2021); ideally, hot spots of belowground biodiversity could serve as a criteria for defining conservation areas.

## Acknowledgments

Fondecyt No. 1190642 (ANID – Chile). To Carlos A. Guerra for helpful suggestions.

## SUPPLEMENTAL FIGURES

**Figure S1.**
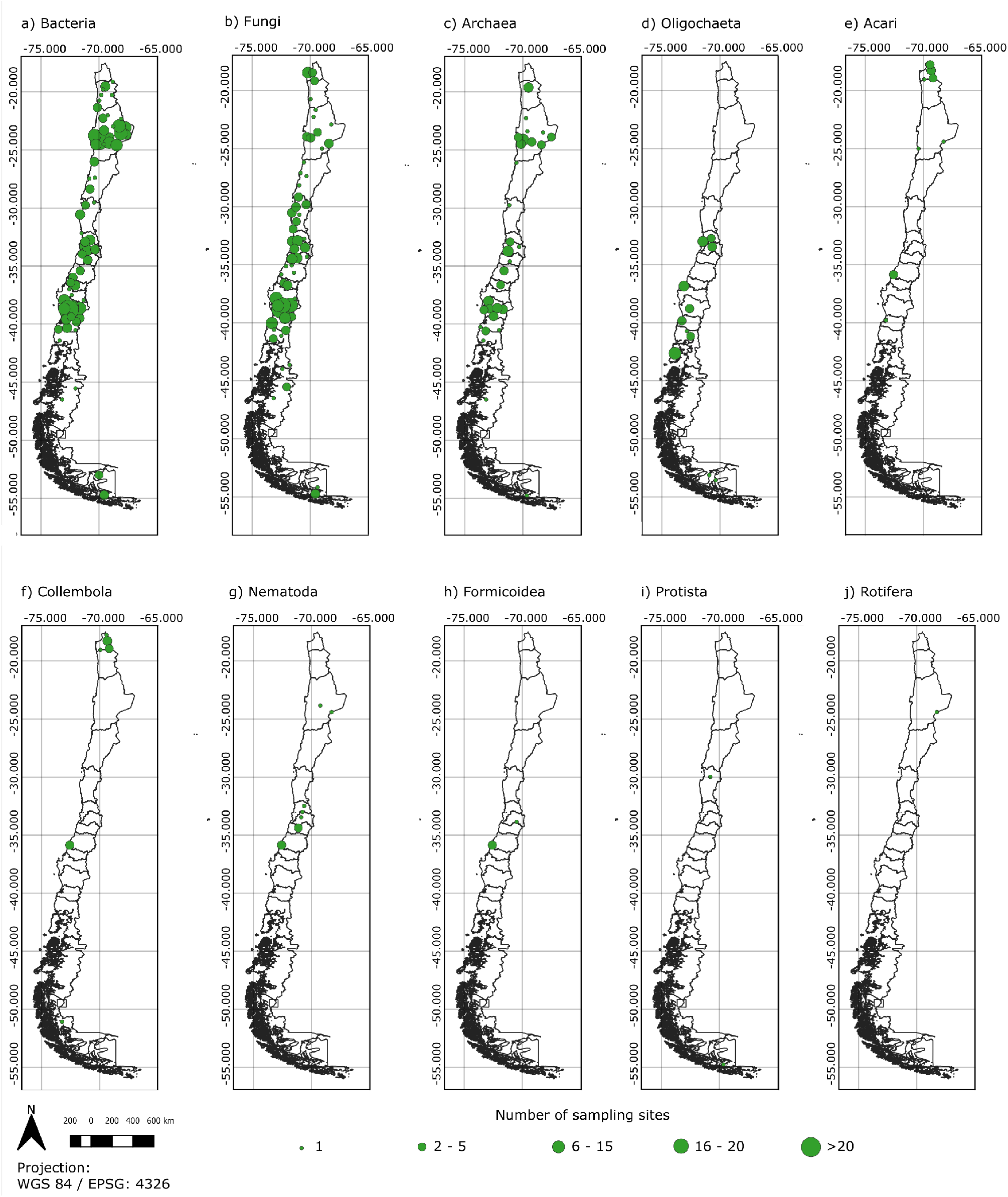
Distribution of the 10 soil taxonomic groups in continental Chile. **a.** Bacteria. **b.** Fungi. **c.** Archaea. **d.** Oligochaeta. **e.** Acari. **f.** Collembola. **g.** Nematoda. **h.** Formicoidea. **i.** Protista. **j.** Rotifera. The size of the circles is based on a 50 km grid.

**Figure S2.**
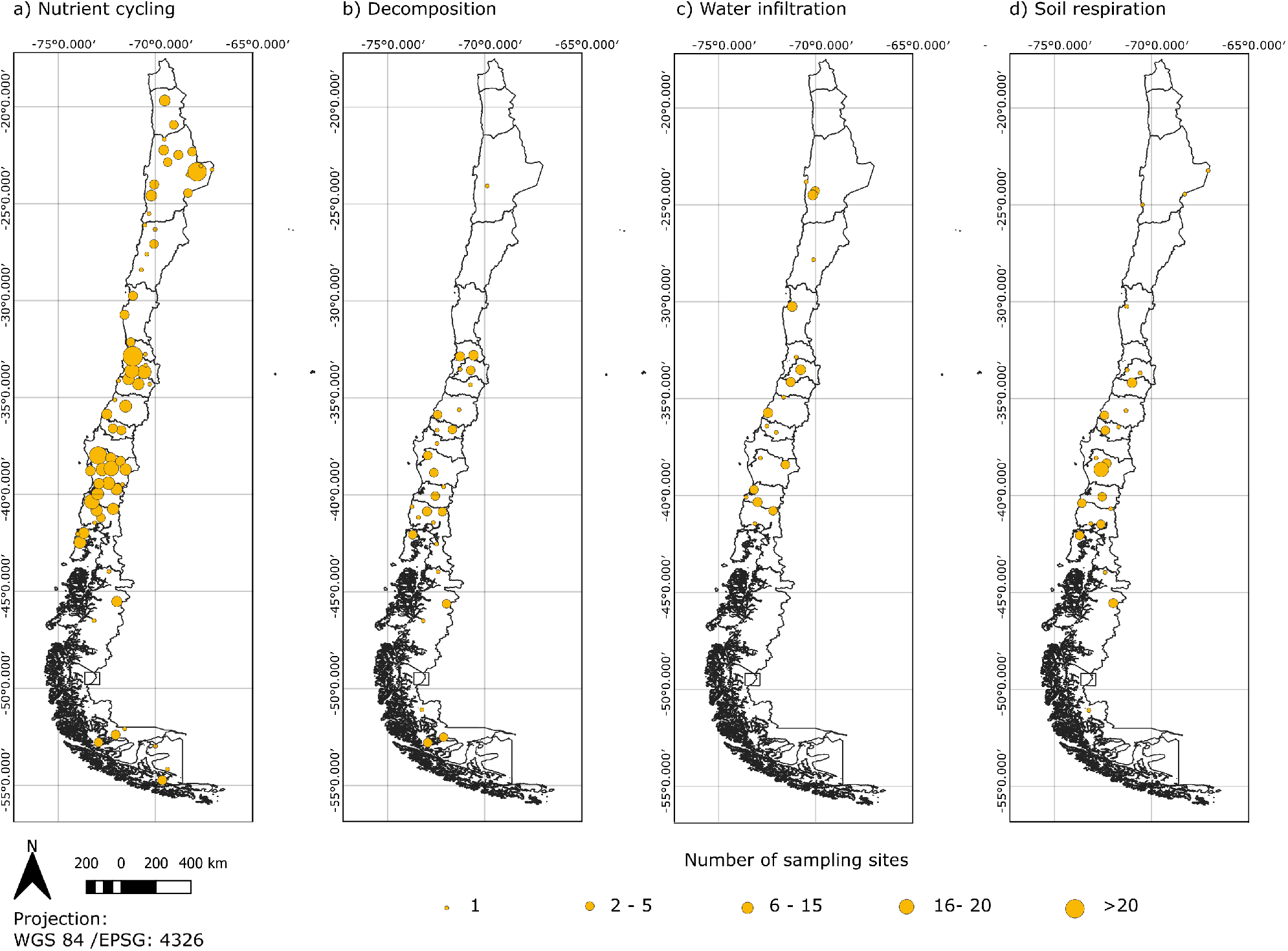
Distribution of the four soil ecosystem functions in continental Chile. **a.** Nutrient cycling. **b.** Decomposition. **c.** Water infiltration. **d.** Soil respiration. The size of the circles is based on a 50 km grid.

